# Loss of HOXA10 causes endometrial hyperplasia progressing to endometrial cancer

**DOI:** 10.1101/2022.04.04.486936

**Authors:** Anuradha Mishra, Nirmalya Ganguli, Subeer S. Majumdar, Deepak Modi

**Author notes:** Corresponding author, Address for Correspondence: Dr. Deepak Modi, Molecular and Cellular Biology Laboratory, National Institute for Research in Reproductive and Child Health, J.M. Street, Parel, Mumbai 400 012, India.

## Abstract

Endometrial cancer is the fourth most common malignancy in women and the precursor lesion is endometrial hyperplasia. HOXA10 is a transcription factor that plays key roles in endometrial functions such as the endowment of receptivity, embryo implantation, and trophoblast invasion. Herein, using testicular transgenesis, we developed transgenic mice that expressed an shRNA against HOXA10 and observed that in these animals there was nearly 70% reduction in the expression of HOXA10. We termed these animals as HOXA10 hypomorphs and observed that downregulation of HOXA10 led to the development of endometrial hyperplasia and most animals developed well-differentiated endometrial adenocarcinoma with age. There was an increased proliferation of the uterine glands and stromal cells in the hypomorphs along with a gain of OVGP1 expression and increased levels of ERα and ERβ. In parallel, there was increased expression of *Wnt4* and β-Catenin, SOX9 and YAP1. We propose that chronic reduction in HOXA10 expression disrupts multiple pathways in the uterus that aids in the development of endometrial hyperplasia which progresses to endometrial cancer with age.

## Introduction

Endometrial cancer is the fourth most common malignancy in women. GLOBOCAN cancer statistics in 2020 estimated that 604127 women globally, and 123907 women in India alone suffered from endometrial cancer (Sung *et al*. 2021). Moreover, the mortality rate in women with endometrial cancer was estimated to be almost 56%. According to the International Agency for Research on Cancer, the incidence rate of endometrial cancer is increasing rapidly and is predicted to increase by more than 50% in the next two decades (Sung *et al*. 2021). Unfortunately, the etiology of endometrial cancer is poorly defined which limits the development of effective diagnostic and therapeutic strategies for this devastating disease.

The precursor lesion for endometrial cancer is endometrial hyperplasia. Endometrial hyperplasia is an abnormal proliferation of the endometrial epithelium and is associated with a significant risk of progression to endometrial cancer. Through the unopposed action of estrogen, the endometrial epithelium proliferates and acquires the malignant phenotype (Huvila *et al*. 2021). Progesterone is known to oppose the action of estrogen and is found to be therapeutic in women with endometrial cancers. However, progestin treatment is effective only in women with atypical complex endometrial hyperplasia and early endometrial carcinoma but the recurrence rate is quite high (Mittermeier *et al*. 2020; Chae-Kim *et al*. 2021). Therefore, it is imperative to identify the factors and mechanisms that govern the progression of endometrial hyperplasia to endometrial cancer.

Several regulatory proteins, growth factors, and their receptors are identified to play a crucial role in endometrial function. Notably, the homeobox gene HOXA10 is well known for uterine development, embryo implantation, and early placentation (Godbole *et al*. 2007; He *et al*. 2018; Ashary *et al*. 2020). During embryonic development, HOXA10 is necessary for uterine biogenesis (Du & Taylor 2016; He *et al*. 2018). Studies show that as compared to fertile women, the expression of HOXA10 is reduced in endometria of women with infertility and recurrent implantation failure (Matsuzaki *et al*. 2009; Cakmak & Taylor 2011; Yang *et al*. 2017). Also, HOXA10 knockout female mice are infertile, the embryos fail to implant due to compromised decidualization. (Benson *et al*. 1996). In endometrial epithelial and stromal cells, HOXA10 expression is regulated by progesterone. Treatment of endometrial stromal or epithelial cells with progesterone upregulates the expression of HOXA10 (Godbole & Modi 2010). Progesterone receptor binding sites are identified on the HOXA10 promoter suggesting that progesterone directly activates the expression of HOXA10 (Yao *et al*. 2003). Gene expression profiling has revealed that several functions of progesterone in the uterus are mediated by HOXA10 and low expression of HOXA10 is observed in endometriosis which is characterized by progesterone resistance (Yao *et al*. 2003; Cakmak *et al*. 2010). These results imply that HOXA10 is essential for mediating some of the functions of progesterone and maintaining the uterus in a differentiated state. Loss of HOXA10 might compromise the differentiated state of the endometrium leading to endometrial disorders. Interestingly, reduced expression of HOXA10 is observed in the endometrial tissue of women with endometrial hyperplasia and endometrial cancer (Matsuzaki *et al*. 2009; Zhong *et al*. 2011). This loss of HOXA10 appears to be due to hypermethylation of the HOXA10 promoter in the endometrial tissue (Fambrini *et al*. 2013). Remarkably, the down-regulation of HOXA10 expression and the methylation in the HOXA10 promoter are strongly correlated with increased tumor grade (Yoshida *et al*. 2006). Furthermore, in vitro studies have demonstrated that forced overexpression of HOXA10 inhibits invasion of endometrial cancer cell lines and tumor dissemination in nude mice (Yoshida *et al*. 2006). These observations imply that HOXA10 might be involved in the pathogenesis of endometrial cancers. However, it is not clear whether the loss of HOXA10 causes endometrial hyperplasia or endometrial cancer.

To understand the functions of genes and their roles in pathological conditions, gene knockout mouse models have been extensively utilized. While the data from these studies are unprecedented and of high value; the knockout mouse model has some inherent issues. Firstly, not all genes can be knocked out successfully; secondly, often the phenotypes are hard to explain due to compensation by related genes (Doyle *et al*. 2013; El-Brolosy & Stainier 2017). Furthermore, the phenotypes of gene knockout mice cannot always be directly extrapolated to the human condition. The expression of the genes in diseased conditions is partially suppressed and not completely lost like what is observed in knockout animals. Thus, to understand the role of a gene in a diseased state, knockdown mouse models are also preferred (Kleinhammer *et al*. 2010; Usmani *et al*. 2013). In the knockdown model, an shRNA against the gene of interest is expressed which allows baseline expression of the gene and protein in the tissues akin to that seen in diseased states.

To comprehend the involvement of HOXA10 in the endometrium and in the causation of endometrial disorders, we developed transgenic mice expressing shRNA against HOXA10 mRNA. Herein, we present the detailed characterization of these mice and show that downregulation of HOXA10 expression in these mice is associated with endometrial hyperplasia that leads to endometrial carcinoma with age.

## Materials and Methods

Institutional Animal Ethics Committee (IAES) approval was taken for the study under projects no, 08/18 and 7/21.

### Generation of Transgenic Animals using Testicular Transgenesis

To identify the shRNA that may lead to effective knockdown for HOXA10 in vivo, two constructs [V3LHS_300885 and V3LHS_300887] that target both human and mouse HOXA10 genes were used (Dharmacon, Colorado, USA). Both these constructs were individually transfected in Ishikawa cells (human endometrial epithelial cells) using Xtreme Gene HP transfection reagent (Sigma Aldrich; Missouri, United States) as per the manufacturer’s protocol. As a control, the non-targeting shRNA [V3LHS_65890S] (Dharmacon) was used. After 72 hours of transfection, the RNA was isolated as described below. The knockdown efficiency was determined by measuring the mRNA levels of *Hoxa10* using real-time PCR.

As the V3LHS_300887 construct gave nearly 70% knockdown efficiency (not shown), this construct was used to generate the transgenic mice. The vector was double digested with restriction enzymes Ssp1 and Pme1 (Fermentas, Massachusetts, US) that released a 7.2 kb fragment **(Fig.1A)**. This linearized 7.2 kb construct contained the necessary elements to drive the transgene expression. It contained Cytomegalovirus CMV promoter, Turbo GFP (tGFP), IRES (Internal ribosomal entry site), Puromycin resistance gene (Puro^R^) along with DNA sequence for transcribing shRNA targetting HOXA10 **(Fig.1A)**.

**Figure 1:**
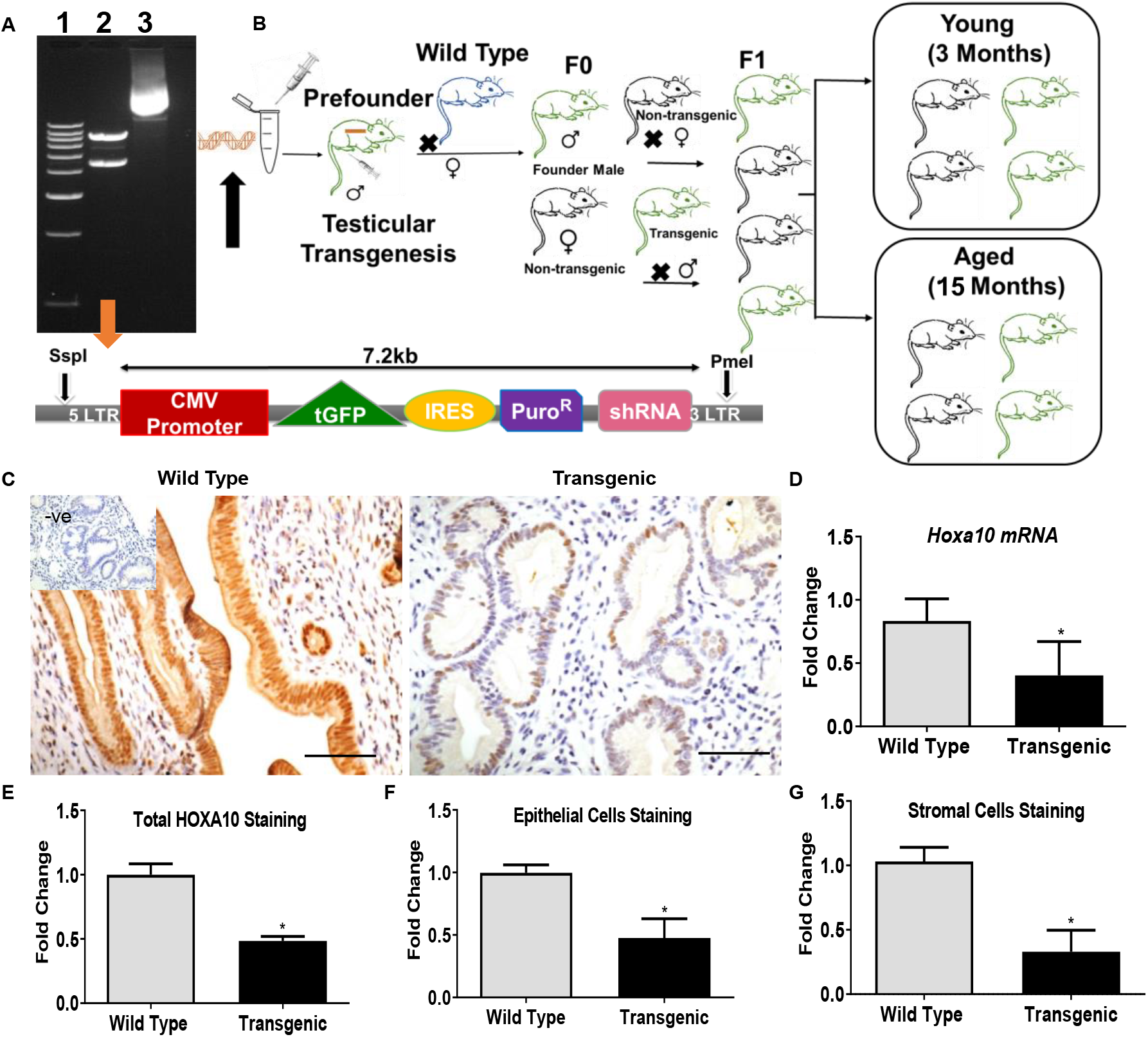
Generation and characterization of *Hoxa10 shRNA* transgenic mice. **A**. Agarose gel electrophoresis showing digested products of pGIPZ plasmid vector; lane 1: ladder (1kb), lane 2: digested plasmid, lane 3: undigested plasmid. The internal elements in the linearized 7.2 kb construct has also shown **B**. Cartoon of the study design where wild type (Blue), Transgenic (Green), and Non-Transgenic (Black) **C**. Immunohistochemistry for HOXA10 in the endometrium of wild type (C57BL/6) and transgenic mice (brown staining). Negative (-ve) section is incubated without primary antibody. Uterine sections are counterstained with hematoxylin (blue staining); Bar, 50μm. **D**. mRNA levels of *Hoxa10* in the endometrium of wild type and transgenic mice. Mean and ±SD for each group is shown (n=3 wild type, n=5 Transgenic). Immunostaining quantification for HOXA10 in the endometrium of wild type and transgenic mice **E**. Total HOXA10, **F**. Epithelial cell stain, **G**. Stromal cell stain. Y-axis represents fold change where the mean levels of wild types are taken as 1. Mean ±SD for each group is shown (n=4 biological replicates per group). Statistically significant differences between the groups are shown by * p≤0.05

After validating the efficiency of this linearized construct (not shown), it was injected in the testis of 30 days old mice as detailed previously (Usmani *et al*. 2013). In brief, two male mice (FVB/J), 30 days old, were anesthetized and hair was shaved off from the lower abdominal and scrotal region. The mice were placed on a piece of autoclaved blotting paper and the scrotal region was wiped with betadine solution. Linearized plasmid DNA (7.2 kb) was injected slowly into both testes of each animal. The testes were held together between a sterile tweezer type electrode and electric pulses were delivered via an electric pulse generator for their simultaneous electroporation. The scrotum was closed and mice were allowed to recover.

Furthermore, these pre-founder males were transported and maintained at the experimental animal facility of ICMR-NIRRCH. Once the males were 8 weeks old, they were mated with wild-type females (C57BL/6). The pups, thus, born were genotyped by PCR as described below. PCR positive founder males were identified and used to develop the colony (**Fig.1B)**.

### Experimental animals

Female transgenic mice were euthanized at 12 weeks (designated as young) of age while some of their siblings were euthanized when they were 15 months of age (designated as aged). The control animals were age-matched females housed in the same conditions.

### Genotyping

Genotyping was done using the Terra PCR Direct Polymerase Mix and kits (Takara Bio; Shiga, Japan). The primers used for genotyping are given in **supplementary table 1**. PCR products were visualized by 2% agarose gel electrophoresis.

### Real-Time PCR (qPCR)

Total RNA was extracted using Trizol reagent (Invitrogen^TM^; Massachusetts, USA), and reverse-transcribed by using high-capacity cDNA reverse transcriptase kit (Applied Biosystems^TM^; Massachusetts, USA) described previously (Mishra *et al*. 2020). The relative mRNA levels of *18S rRNA, Hoxa10, Esr1, Esr2, Wnt4*, and *Ctnnb1* were estimated by qPCR using iQ SYBR green chemistry (Bio-Rad; CA, USA). The primer sequences for the same are given in **supplementary table 1**.

Quantitative PCR was carried out in the CFX-96 thermal cycler (Bio-Rad). All PCR reactions were carried out in duplicates, gene expression was normalized to the levels of *18S*, and fold change was calculated using the Pfaffl method as described earlier (Mishra *et al*. 2020).

### Histological analysis

Five micrometers thick paraffin sections of 4% PFA fixed tissues were cut and mounted on poly-L-lysine (Sigma Aldrich) coated slides. Sections were deparaffinized in xylene, hydrated in descending grades of alcohol, stained with hematoxylin and eosin (Himedia; Maharashtra, India), and mounted using DPX mountant. Slides were viewed under a bright field microscope (Olympus; Tokyo, Japan) and representative areas were photographed.

### Immunohistochemistry

To determine the abundance and cellular distribution of HOXA10, Cytokeratin, OVGP1, MKi-67, ERα, ERβ, SOX9, β-catenin, and YAP1, immunohistochemistry was performed as described earlier (Mishra *et al*. 2020). Briefly, the antigens were retrieved by boiling in Tris-EDTA Buffer, pH9, or Sodium Citrate Buffer, pH 6. Blocking was done for 1 hour in 5% BSA (MP Biomedicals; Maharashtra, India) and for HOXA10, 1% donkey serum (Jackson Immunology; Pennsylvania, United States) was used. Sections were probed overnight with a primary antibody. Negative controls were incubated with either the isotype IgG or PBS instead of primary antibodies. The next day, slides were washed and incubated in a biotinylated secondary antibody followed by streptavidin-HRP (ABC kit Santa Cruz Biotechnology; Texas, USA). Detection was done using hydrogen peroxidase as substrate and 3, 3’-diaminobenzidine (DAB) (Sigma Aldrich) as a chromogen. All sections were counterstained with hematoxylin and mounted in DPX. Slides were viewed under a bright-field microscope and representative areas were photographed (Olympus). The sources and the optimized concentrations of primary antibodies are given in **supplementary table 2**.

### Quantification of Immunostaining

“Fiji” version of ImageJ software was used to quantify the intensity of the chromogen and gland to stroma ratio in uterine sections (Schindelin *et al*. 2012). Along with total intensity, for MKi-67, SOX9, β-catenin, and YAP1, the number of positive nuclei was also estimated using the “Qupath v0.2.3” (Bankhead *et al*. 2017). At least three random sections were analyzed per animal and the mean value was obtained to estimate the level of expression. To quantify cell type changes in immunostaining, five different areas of stroma and epithelia were selected per section per animal and quantified as above.

### Statistical Analysis

Experimental data was calculated in terms of mean and standard deviation. All the statistical analysis was done using two-way repeated-measures ANOVA and Mann–Whitney test or unpaired Student’s t-test in GraphPad Prism (Version 8; California, USA). Results with p ≤ 0.05 were accepted as statistically significant.

## Results

### The expression of HOXA10 is downregulated in the endometrium of transgenic mice

To determine the abundance and localization of HOXA10, immunohistochemistry was performed (n=4 biological replicates per group). As shown in **Fig.1C**, HOXA10 was localized to the nucleus and cytoplasm of glandular epithelium and stromal cells in the endometrium of the wild-type mice. In transgenic mice, expression of HOXA10 was reduced when compared to the wild-type controls. Quantitatively, almost 50% of reduction in the expression of HOXA10 was observed in transgenic mice as compared to wild type and this reduction was statistically significant (P-value≤0.05). Also, there was a significant reduction in the expression of HOXA10 in both epithelial and stromal cells of transgenic mice as compared to the wild-type female mice **(Fig.1E-1G)**.

Real-time PCR was also performed to test the mRNA levels of HOXA10 in the diestrus stage uteri of transgenic female mice (n=5). As compared to the wild-type controls (n=3), there was more than a 50% reduction in the mRNA levels of *Hoxa10* in the uteri of transgenic mice **(Fig.1D)**.

Since the expression of HOXA10 was not completely switched off but was downregulated by shRNA, we termed these animals as HOXA10 hypomorphs and this term is used throughout the manuscript.

### HOXA10 hypomorphs develop endometrial hyperplasia that progresses to endometrial cancer

Gross morphology of the uterine horns from young hypomorphs appeared almost similar to their age-matched wild-type controls **(Fig. 2A)**. However, as compared to the controls, in the aged hypomorphs, the uteri appeared thicker and often had nodule-like structures **(Fig 2A)**. In one aged hypomorph, a large mass was observed in the uterus that appeared like a tumor **(Supp. Fig.1)**.

**Figure 2:**
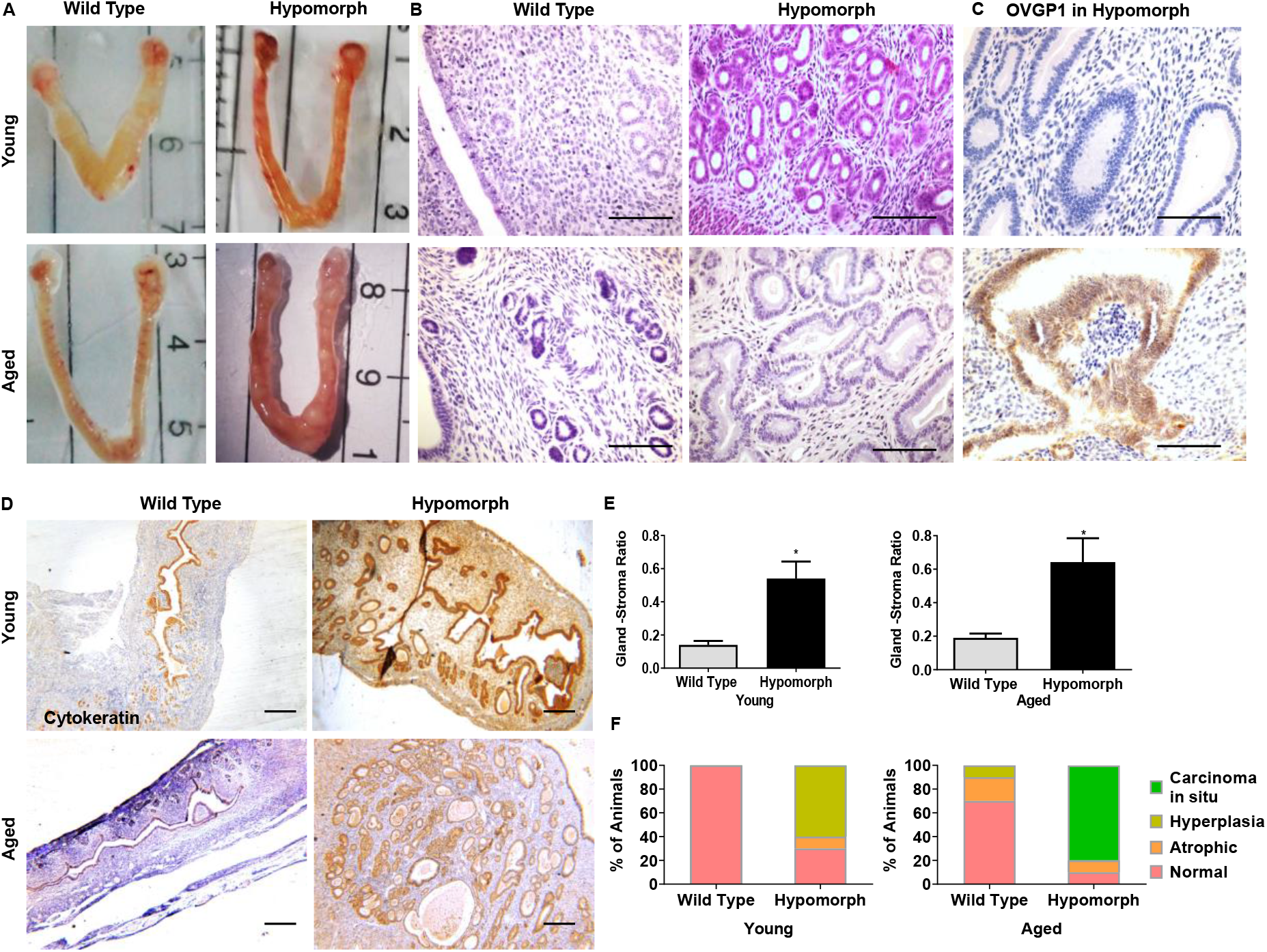
Endometrial hyperplasia progresses to endometrial cancer in mice hypomorphic for HOXA10. **(A)** Representative images of young (3 months) and aged (15 months) HOXA10 hypomorphic female mice uterus **(B)** Hematoxylin and Eosin stained endometrium of wild type, and hypomorphic female mice (n=10 biological replicates per group); Bar, 50μm. **(C)** Immunohistochemistry for OVGP1 in young and aged hypomorphs **(D)** Immunostaining of cytokeratin in endometrial tissues of young and aged wild type and hypomorphic mice (Brown staining). Uterine sections are counterstained with hematoxylin (blue staining). Negative (-ve) control section is incubated without primary antibody. Bar, 50μm for OVGP1 and bar, 100μm for Cytokeratin (**E**) Gland-stroma ratio in the endometrium of young and aged hypomorphs as compared to wild types in diestrus phase. Mean ±SD is shown for n=5 wild types, n=5 hypomorphs. Statistically significant differences between the groups are shown by * p≤0.05 **(F)** Graphs representing the proportion of mice with normal, atrophic endometrium, uterine hyperplasia, and carcinoma in situ of endometrium in wild type and hypomorphs (n=10 biological replicates per group).

Histologically, as compared to the age-matched wild-type females (n=10), the endometrium of young hypomorphs (n=10) had large number of glands **(Fig.2B)**. There was a wide variation in gland size, many had cystic dilatation or irregular luminal contours **(Fig.2B)**. Cytological atypia if any was rare. As the animals aged, in most controls (n=10), the uterine histology appeared normal, although, the glands appeared smaller and few in numbers. In a proportion of animals, the uterine tissue appeared atrophied. In the aged hypomorphs (n=10), there were changes resembling endometrial adenocarcinoma. There were small round multiple back-to-back glands with loss of intergranular stroma **(Fig 2B, Supp. Fig.2A&2B**), there was glandular bridging and had taken the cribriform appearance, in some areas, there was villoglandular appearance **(Supp. Fig.2C&2D**). The cell polarity was largely preserved and the epithelium appeared pseudostratified. In some instances, the nuclei appeared round and there was a loss of cell polarity in the epithelium. The stroma appeared edematous with a large number of inflammatory cells. In two aged hypomorphs, the stroma appeared desmoplastic (not shown). In both young and aged hypomorphs, the gland to stroma ratio was almost 4 times that in the controls and the increase was statistically significant (p≤0.05, **Fig.2E**). Further, in the young hypomorphs, 60% of the animals had developed uterine hyperplasia. In one animal, the uterus appeared atrophied while in others, the histology of the endometrium was normal **(Fig.2F)**. In the aged controls, 70% of animals had a normal uterus and in 20% of animals, the uterus had atrophied endometrium while only in one animal endometrial hyperplasia was observed. In the aged hypomorphs, one animal had normal a uterus and one had an atrophied endometrium. In the remaining animals, there were changes resembling endometrial cancer with concomitantly uterine hyperplasia **(Fig.2F)**.

OVGP1 is a specific marker expressed only in endometrial cancer (Wang *et al*. 2010). Immunohistochemistry was performed to localize the expression of OVGP1 in endometrial tissues of young and aged hypomorphs and their age-matched wild-type controls (n=3/group). We did not observe the expression of OVGP1 in wild-type uteri of control animals and young hypomorphs. Cytoplasmic expression of OVGP1 was observed only in the glandular epithelium of aged hypomorphs **(Fig.2C)**.

Immunohistochemistry using cytokeratin revealed a marked increase in the number of uterine glands in *Hoxa10* hypomorphs as compared to their age-matched wild-type controls (n=3/group) as shown in **Fig.2D**. The number of glands increased further and the glandular organization and complexity were higher in aged hypomorphs but the myometrial invasion was not observed in any of the animals.

### Cell proliferation in uteri of HOXA10 hypomorph

As shown in **Fig.3**, irrespective of the age (n=3/group), a few cells were positive for MKi-67 in wild-type uterus, large numbers of epithelial and stromal cells were positive in hypomorphs **(Fig.3A)**. Quantitative analysis shows that less than 10% of cells were positive for MKi-67 in both young and aged wild types while ~ 20% of cells were positive for MKi-67 in young hypomorphs and ~ 35% of cells in aged hypomorphs. This upregulation was statistically significant (p≤0.001, **Fig.3B**)

**Figure 3:**
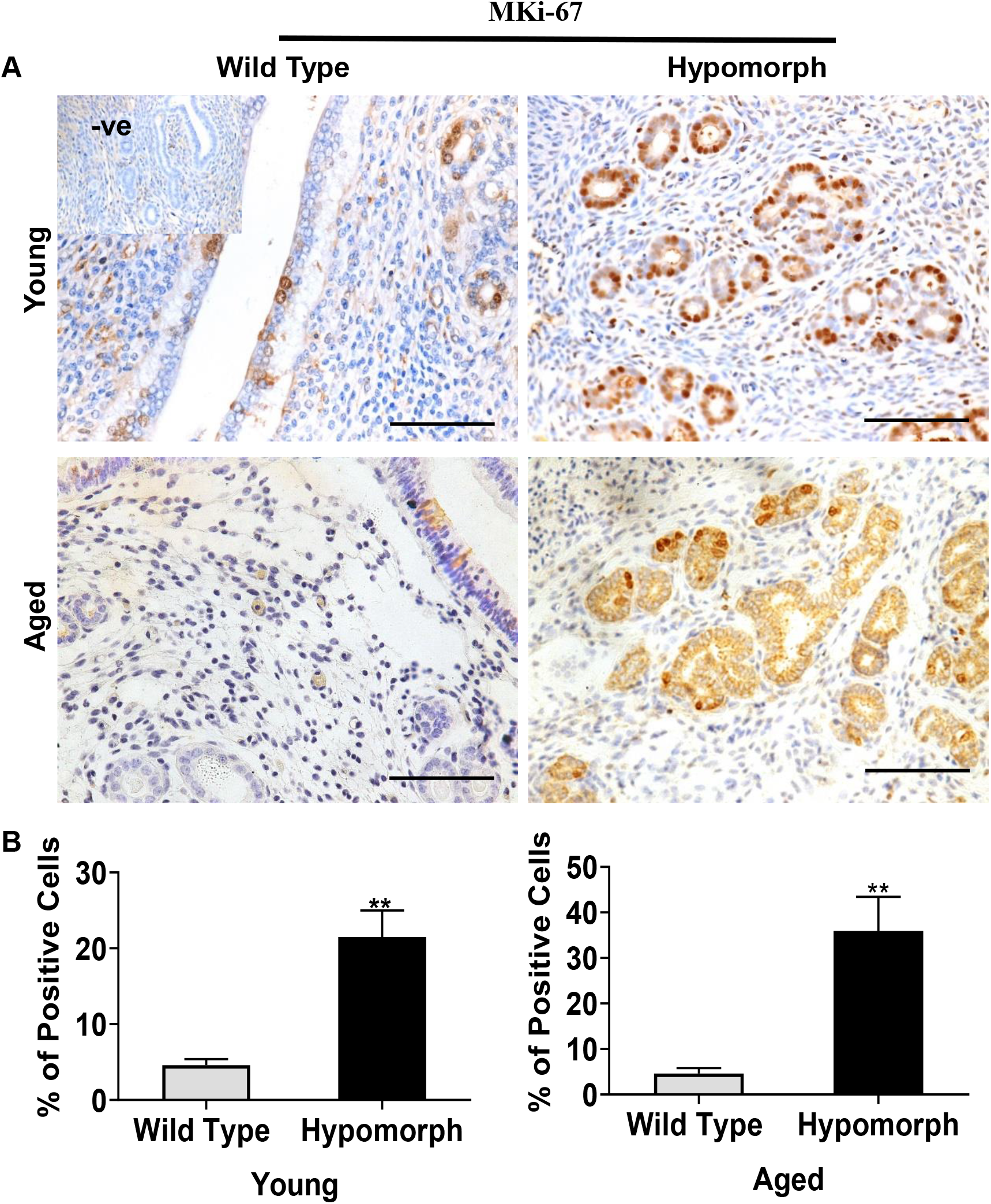
Proliferation in the endometrium of HOXA10 hypomorphs. **(A)** Immunohistochemistry for MKi-67 antibody in the endometrium of young (3 months) and aged (15 months) wild types and hypomorphs (Brown staining). Uterine sections are counterstained with hematoxylin (blue staining). Negative control is section incubated without primary antibody. Bar, 50μm. **(B)** Immunostaining quantification where Y-axis represents the percentage of MKi-67 positive cells. Mean and ±SD for each group is shown (n=3 biological replicates per group). Statistically significant differences between the groups are shown by ** p≤0.001

### Loss of HOXA10 alters the expression of estrogen receptors in HOXA10 hypomorphs

The expression of ERα appeared almost similar in the uteri of young hypomorphic animals (n=3) as wild type controls (n=3), this was mainly in the glandular epithelium (**Fig.4A&4C)**. In the aged hypomorphs (n=3), ERα expression was high in the glands and stroma. There was a ~2 fold increase in the ERα protein levels in the aged hypomorphs as compared to wild-type controls (n=3) and this increase was statistically significant (p≤0.00001) (**Fig.4C)**.

**Figure 4:**
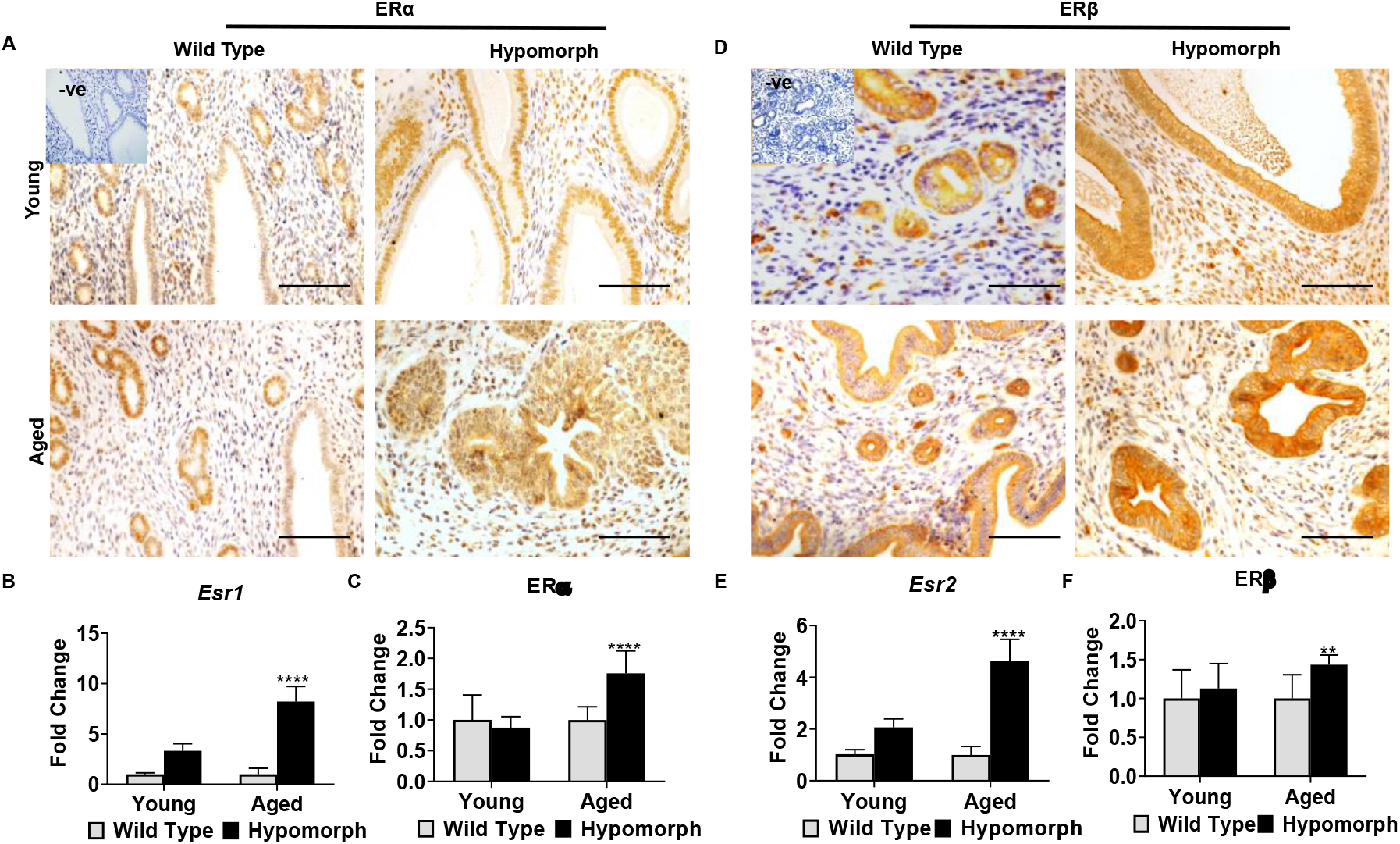
Expression of estrogen receptors in the endometrium of HOXA10 hypomorph. **(A&D)** Immunohistochemistry for ERα, and ERβ in the endometrium of young (3 months) and aged (15 months) wild type, and hypomorphic mice (Brown staining). Uterine sections are counterstained with hematoxylin (blue staining). Negative (-ve) control section is incubated without primary antibody. Bar, 50μm. **(C&F)** Quantification of ERα and ERβ protein in the endometrium of hypomorphic female mice. Y-axis represents the fold change where the mean level of wild types are taken as 1. Mean and ±SD for each group is shown (n=3/group) **(B&E)** qPCR for estrogen receptors *Esr1*, and *Esr2*, in the endometrium of hypomorphic female mice and their age-matched controls. Y-axis is showing fold change where mean level of wild types are taken as 1. Mean ±SD for each group is shown (n=3 biological replicates per group). Statistically significant differences between the groups are shown by ** p≤0.001, **** p≤0.00001

As compared to the age-matched controls, there was an increase in *Esr1* mRNA levels in the young and aged hypomorphs. In the young hypomorphs as compared to controls, the increase in *Esr1* mRNA levels was almost 3 folds while in the aged hypomorphs, the increase in the expression of *Esr1* was almost ~8 folds and this increase was statistically significant (p≤0.00001, **Fig.4B**).

As compared to controls, there was an increase in the expression of ERβ in the hypomorphs irrespective of age (**Fig.4D)**. Quantitatively, ERβ levels were marginally increased in young hypomorphs as compared to wild-type controls; however, this upregulation was not statistically significant. In the aged hypomorphs, ERβ levels were significantly increased (p≤0.001) by ~1.5 folds as compared to their age-matched wild-type controls **(Fig. 4F)**.

In comparison to age-matched controls, in young animals, there was a ~2 folds increase in the expression of *Esr2* and ~5 folds increase in aged hypomorphs and this increase was statistically significant in aged hypomorphs (p≤0.00001, **Fig.4E)**.

### Altered expression of β-catenin in the uterus of HOXA10 hypomorphs

Immunohistochemical analysis was performed to localize β-catenin in the endometrial tissues of young and aged hypomorphs and their age-matched wild-type controls (n=3/group) as shown in **Fig.5**.

**Figure 5:**
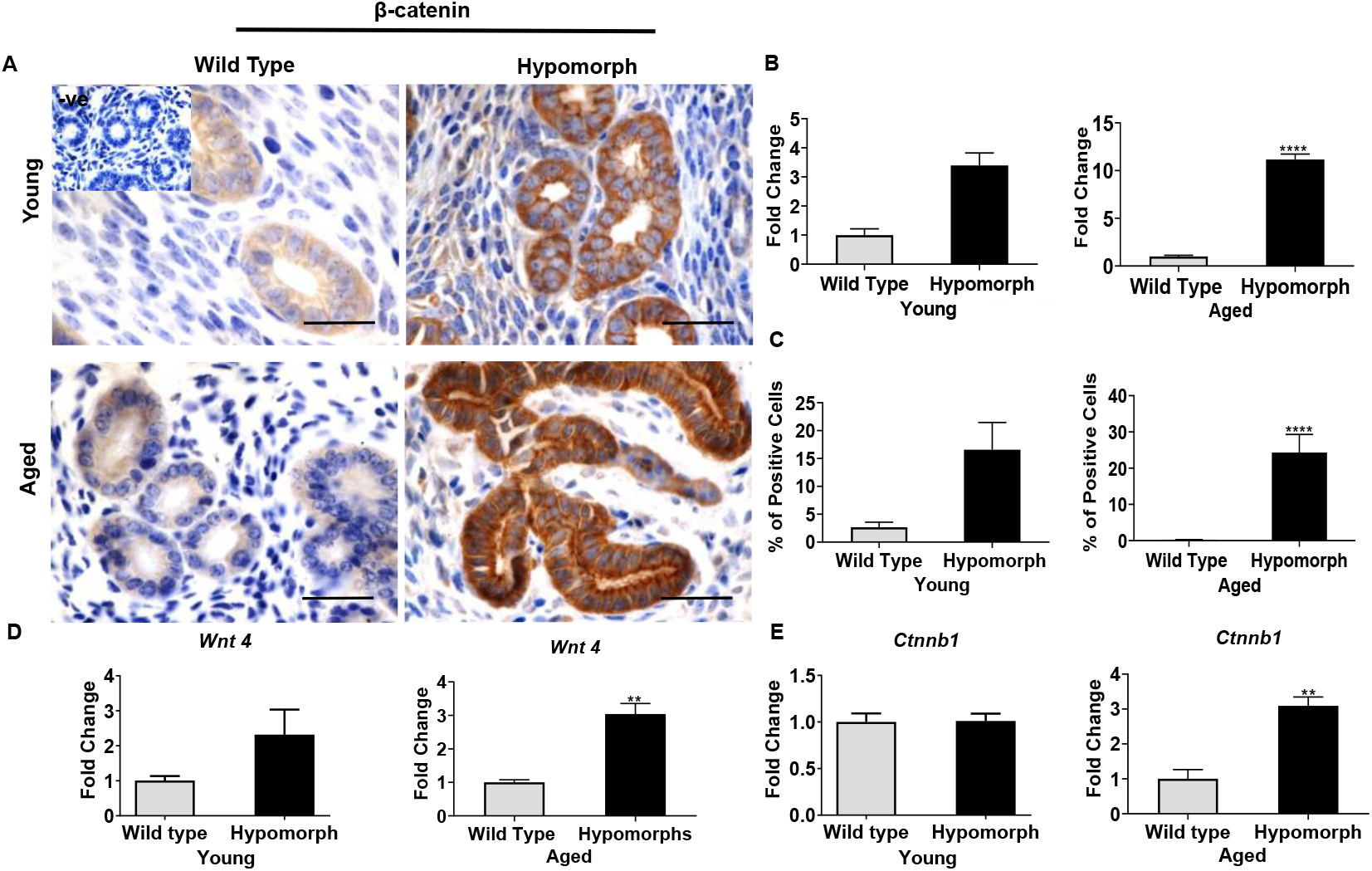
Expression of β-catenin and Wnt4 in the endometrium of HOXA10 hypomorph. **(A)** Immunohistochemistry for β-catenin protein in endometrial tissues of young (3 months) and aged (15 months) wild type and hypomorphic mice (Brown staining). Uterine sections are counterstained with hematoxylin (blue staining). Negative (-ve) control section is incubated without primary antibody. Bar, 20μm **(B)** Immunostaining quantification shows the protein intensity level of β-catenin where Y-axis is showing the fold change where the mean level of wild types are taken as 1. Mean and ±SD for each group is shown (n=3 wild types, n=3 hypomorphs) **(C)** Here, graphs represent the percentage of nuclear positive cells for β-catenin protein. Mean and ±SD for each group is shown (n=3 biological replicates per group) **(D&E)** qPCR for *Wnt4*, and *Ctnnb1* in the endometrium of hypomorphic female mice and their age-matched controls. Y-axis is showing the fold change where mean level of wild types are taken as 1. Mean ±SD for each group is shown (n=3/group). Statistically significant differences between the groups are shown by ** p≤0.001, **** p≤0.00001

In wild-type females, β-catenin was generally cytoplasmic in the glandular epithelium of endometrial tissue. In hypomorphs, expression of β-catenin was observed in both the nucleus and cytoplasm of the glandular epithelium **(Fig.5A)**. Quantitatively, in the young animals, the expression of β-catenin was ~3 folds higher in the young hypomorphs and ~11 folds in aged hypomorphs and this increase was statistically significant in aged hypomorphs (p≤0.00001, **Fig.5B)**. Since β-catenin can translocate to the nucleus, we quantified the percentage of cells that were positive for nuclear β-catenin. Less than 5% of cells were positive for nuclear β-catenin in wild-type uteri in both young and aged animals. However, this increased to ~15% in young and ~25% in aged hypomorphs. In aged hypomorphs, this increase was statistically significant (p≤0.00001, **Fig.5C**).

There was a marginal increase in *Ctnnb1 mRNA* levels in young hypomorphs while ~ 3 folds significant (p≤0.001) increase in aged hypomorphs when compared with their age-matched wild-type controls (**Fig.5E**). *Wnt4* mRNA expression was also elevated in the uteri of both young and aged HOXA10 hypomorphs. As compared to wild-type controls, there was a ~2 folds increase in *Wnt4* mRNA levels in young hypomorphs and ~3 folds significant (p≤0.001) increase in the aged hypomorphic mice (**Fig.5D**).

### Expression of SOX9 in HOXA10 hypomorphs

SOX9 expression was detected in the uterine glandular epithelium of wild-type controls (n=3/group), the expression, in general, was weak in the wild-type young and aged controls (**Fig. 6A)**. In the young and aged hypomorphs (n=3/group), a large number of cells were intensely stained for SOX9, there was a ~3 folds increase in the expression of SOX9 in the young animals and ~4 folds increase in aged hypomorphs, the increase was statistically significant (p≤0.05, **Fig.6B)**. We quantified the numbers of cells with nuclear SOX9 and observed that ~3% of cells were positive for SOX9 in young wild-type uteri, the number significantly (p≤0.001) increased to 13% in the uteri of young hypomorphs. Similarly, ~2% of cells were positive for SOX9 in aged wild-type uteri, this increased to ~25% in the hypomorphs. This increase was statistically significant (p≤0.00001, **Fig.6C**).

**Figure 6:**
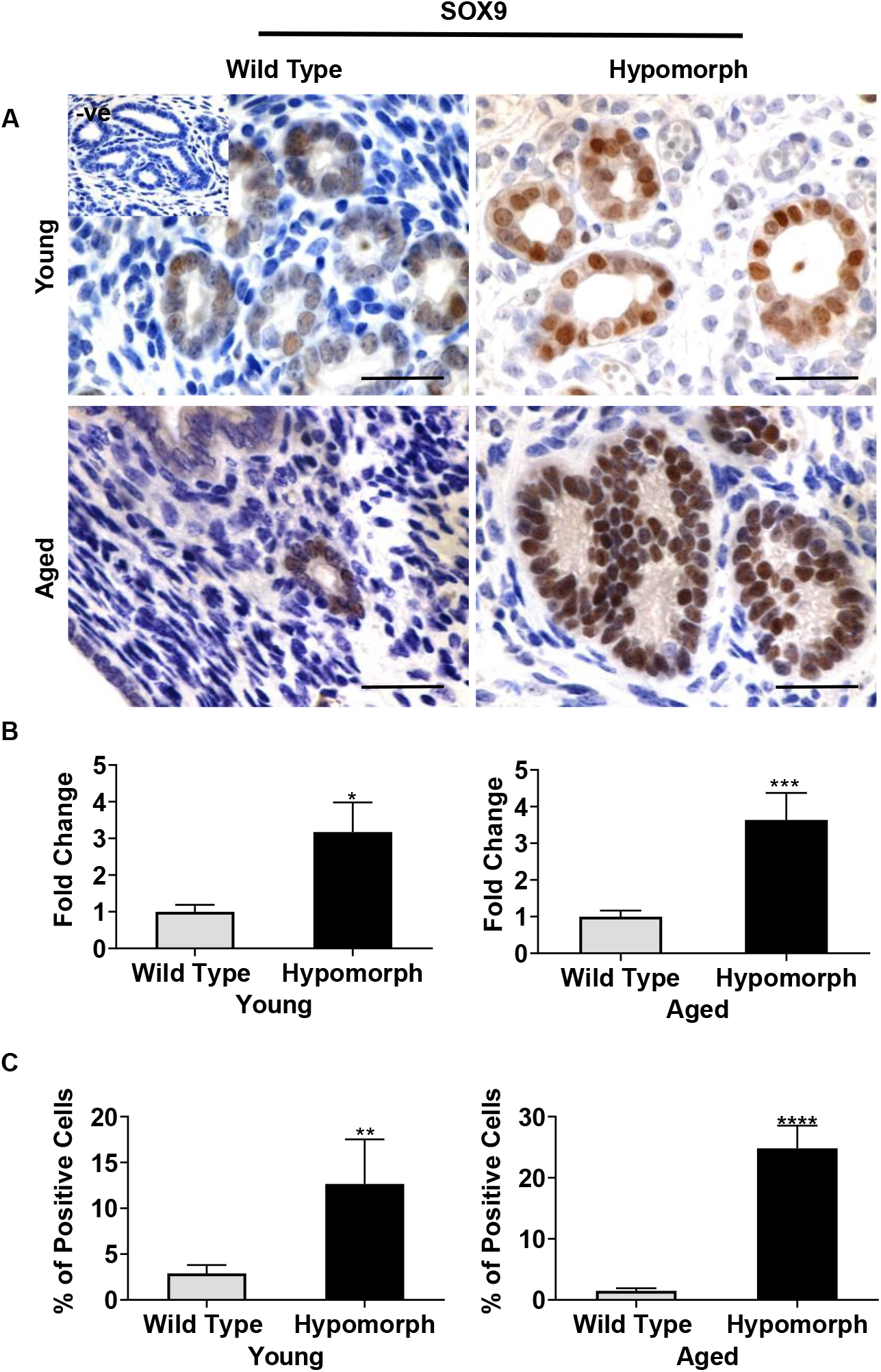
Expression of SOX9 in endometrium of HOXA10 hypomorphs. **(A)** Immunohistochemistry for SOX9 in endometrial tissues of young (3 months) and aged (15 months) wild type and hypomorphic mice (Brown staining). Uterine sections are counterstained with hematoxylin (blue staining). Negative (-ve) control section is incubated without primary antibody. Bar, 20μm **(B)** Immunostaining quantification shows the protein intensity level of SOX9 where Y-axis is showing the fold change where the mean level of wild types are taken as 1. Mean and ±SD for each group is shown (n=3 biological replicates per group) **(C)** Here, graphs represent the percentage of nuclear positive cells for SOX9 protein. Mean and ±SD for each group is shown (n=3 biological replicates per group). Statistically significant differences between the groups are shown by * p≤0.05,** p≤0.001, *** p≤0.0001, ****p≤0.00001

### Expression of YAP 1 in HOXA10 hypomorphs

In wild-type uteri (n=3), YAP1 expression was largely cytoplasmic and restricted to the glandular epithelium. In the young hypomorphs (n=3), YAP1 was largely cytoplasmic; however, the intensity of staining was much higher than that observed in controls **(Fig.7A)**. In the aged controls (n=3), both nuclear and cytoplasmic localization of YAP1 was observed, albeit the expression was weak. In the aged hypomorphs (n=3), intense staining for YAP1 was detected in almost all the glands, the staining was both nuclear and cytoplasmic (**Fig.7A)**. There was a significant (p≤0.05) increase in the expression of YAP1 in both young and aged hypomorphs **(Fig.7B)**. There were ~17% of positive cells for YAP1 in young wild-type uteri, this increased to ~35% in young hypomorphs and this increase was also statistically significant (p≤0.0001, **Fig.7C)**. Similarly, in the aged wild type uteri, ~25% of the cells were positive for YAP1, this significantly (p≤0.001) increased to ~50% in aged hypomorphs (**Fig.7C)**.

**Figure 7:**
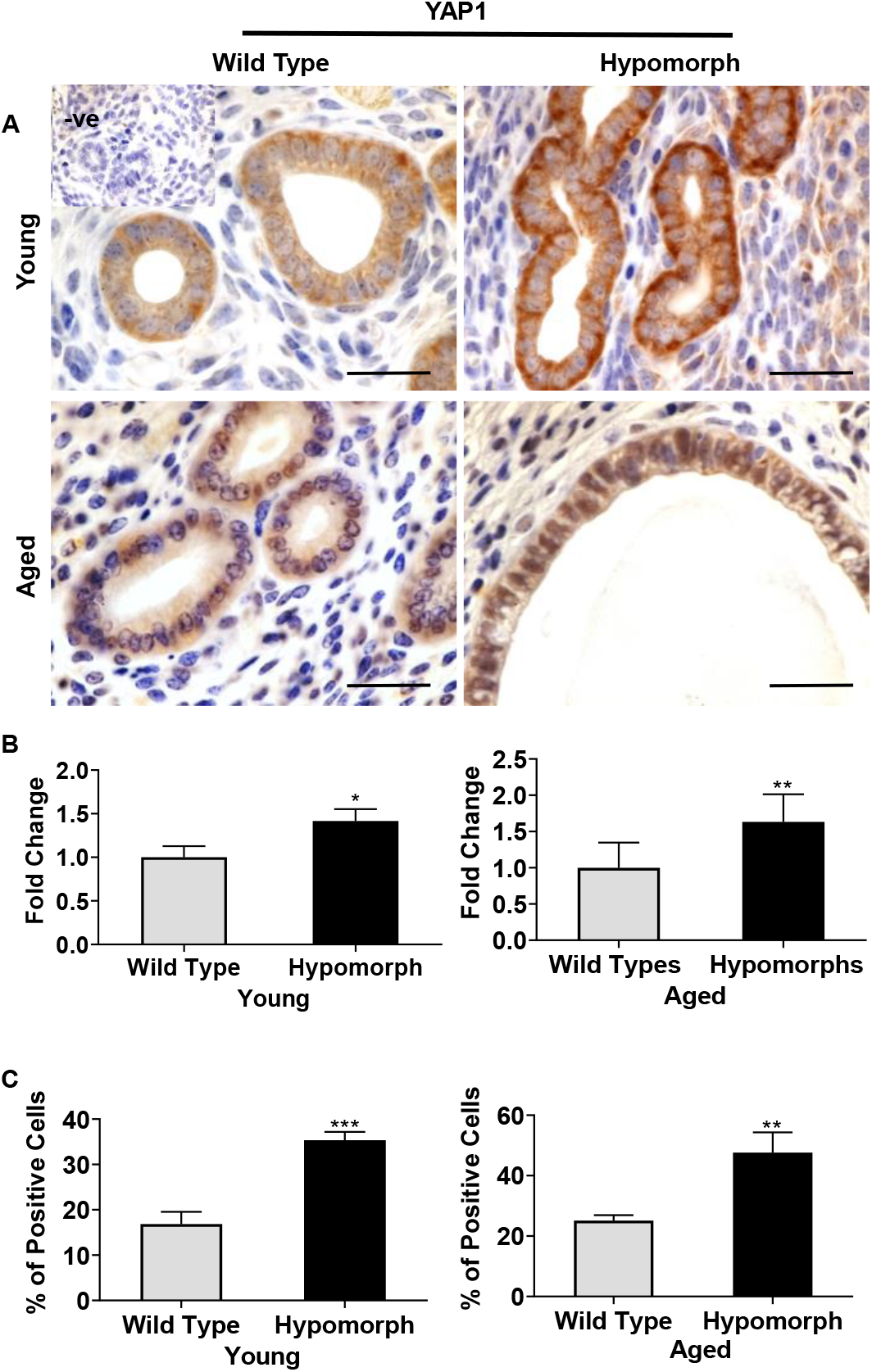
Expression of YAP1 in endometrium of HOXA10 hypomorphs. **(A)** Immunohistochemistry for YAP1 in endometrial tissues of young (3 months) and aged (15 months) wild type and hypomorphic mice (Brown staining). Uterine sections are counterstained with hematoxylin (blue staining). Negative (-ve) control section is incubated without primary antibody. Bar, 20μm **(B)** Immunostaining quantification shows the protein intensity level of YAP1 where Y-axis is showing the fold change where the mean level of wild types are taken as 1. Mean and ±SD for each group is shown (n=3 wild types, n=3 hypomorphs). **(C)** Here, graphs represent the percentage of nuclear positive cells for YAP1 protein. Mean and ±SD for each group is shown (n=3 biological replicates per group). Statistically significant differences between the groups are shown by *p≤0.05, **p≤0.001, *** p≤0.0001

## Discussion

In this study, we developed a transgenic mouse model overexpressing a *Hoxa10* shRNA and studied their uterine phenotypes. Mice with reduced HOXA10 developed endometrial hyperplasia which progressed to an endometrial cancer-like phenotype as they aged. This was associated with increased cell proliferation, and altered expression of estrogen receptors, β-catenin, SOX9, and YAP1.

Endometrial hyperplasia is a precursor lesion to endometrial cancer. However, the molecular mechanisms of endometrial carcinoma development, progression, and invasion/migration have not been identified. Herein, we observed that downregulation of HOXA10 led to the development of an endometrial hyperplasia-like phenotype that progressed to an endometrial cancer-like phenotype with age. OVGP1 is a protein specifically secreted by the oviduct and it is not expressed by the endometrium except during a very narrow window of the embryo implantation (Laheri *et al*. 2018). However, OVGP1 expression is activated in the epithelial cells of 95% of patients with atypical endometrial hyperplasia and most strongly in grade 1 endometrial cancers making it a specific marker for diagnosis of early endometrial cancer (Woo *et al*. 2004). Interestingly, we found that OVGP1 was not expressed in the endometrial epithelial cells in young transgenic animals with hyperplastic endometrium but was detected in the endometrial epithelial cells of aged animals that had cancer phenotype. These observations confirm that chronic loss of HOXA10 promotes the development of low-grade, well-differentiated endometrioid carcinoma in situ. Misexpression of HOX genes including HOXA10 is reported in esophageal squamous cell carcinoma, breast cancer, neuroblastoma, lung cancer, melanoma, bone cancer, blood cancer, colorectal cancer, prostate, ovarian, and cervical cancers (Hajirnis & Mishra 2021; Ponrathnam *et al*. 2021). Interestingly, forced expression of HOXA10 causes endometrioid-like differentiation of mouse ovarian surface epithelium, and overexpression of HOXA10 is seen in ovarian endometrioid adenocarcinoma (Cheng *et al*. 2005; Tanwar *et al*. 2013). However, in endometrial cancer, there is a reduction in HOXA10 expression (Yoshida *et al*. 2006). Thus, it appears that changing levels of HOXA10 gene expression can contribute to cancer in a context-dependent manner.

Homeobox genes are regulators of cell proliferation in multiple tissues and their altered expression is associated with tumor grade (Li *et al*. 2019). Herein, we observed that the numbers of Ki-67 positive cells were elevated in the hyperplastic endometrium of young hypomorphs which further increased in the aged hypomorphs. This implies that loss of HOXA10 causes sustained proliferation of endometrial epithelium. Loss of HOXA10 inhibits cell proliferation in oral squamous cell carcinoma and hepatocellular carcinoma; however, it promotes cell proliferation in testicular cancer and gastric cancer (Chen *et al*. 2018; Song *et al*. 2019; Zhang *et al*. 2019). Although in poorly differentiated endometrial carcinoma cell lines, overexpression of HOXA10 does not alter the rate of proliferation (Yoshida *et al*. 2006), the loss of HOXA10 in a well-differentiated endometrial cancer cell line (Ishikawa) led to an increase in proliferation (our unpublished data). Thus, the effects of HOXA10 on cell proliferation are cell type and context-dependent. Nevertheless, our results demonstrate that in the endometrial epithelium, HOXA10 is required to keep cell proliferation in check and its loss leads to endometrial hyperplasia and endometrial cancer with age.

Estrogen is a key factor required for the proliferation of the endometrial epithelium and prolonged unopposed exposure of animals to estrogen causes endometrial hyperplasia (Goad *et al*. 2018). Estrogen acts via its two classical receptors, ERα and ERβ. Of these ERα is responsible for cell proliferation in the endometrium while ERβ isoforms restrain the ERα-mediated cell-specific mitotic responses (Hapangama *et al*. 2015). We tested the expression of both these receptors and observed that in the young hypomorphs with hyperplastic endometrium, the expression of both ERα and ERβ were unchanged when compared to normal endometrium. However, as the animals aged, the expression of ERα and ERβ was higher in the hypomorphs. This is in contrast to that observed in human endometrial cancers where expression of both these isoforms is reduced (Chakravarty *et al*. 2008; Jarzabek *et al*. 2013). However, intactness of estrogen signaling and normal ER activity is shown to be essential for the proliferation and metastasis of endometrial cancers and the knockdown of ERβ in endometrial cancer cells promotes cell proliferation (Jeong *et al*. 2009; Monsivais *et al*. 2019; Treeck *et al*. 2019). Based on our work, it is reasonable to propose that in absence of HOXA10, the hyperplasia is not due to elevated expression of estrogen receptors; however, there is sustained estrogen receptor expression in absence of HOXA10 with age which may contribute to endometrial cancer.

Endometrial hyperplasia and endometrial cancers are Wnt-dependent conditions where the Wnt pathway is upregulated (Goad *et al*. 2018; Fatima *et al*. 2021). Herein, we observed that in the normal endometrium, *Wnt4* is expressed in the endometrium of young animals which is extinguished with age; however, it is overexpressed in the young HOXA10 hypomorphs and is persistently elevated as the animals’ age. We also observed that in the young hypomorphs, the Wnt downstream target gene, *Ctnnb1*, and its protein product β-catenin were elevated and its expression was further increased in aged animals. The proliferative action of β-catenin is by the virtue of its transcription factor activity, where in response to Wnt signaling β-catenin translocates from the membrane into the nucleus (MacDonald *et al*. 2009). We observed that in the young hypomorphs, the number of cells with nuclear localization of β-catenin was higher than in age-matched controls while in the aged hypomorphs, this number was increased by almost 30 folds. These results indicated that loss of HOXA10 activates the Wnt-β catenin pathway in the endometrium and the expression is sustained in the uteri of aged animals that develop cancer.

The sex-determining region Y box 9 (SOX9) is a transcription factor that plays a key role in developmental processes and is suggested to also act as an oncogene and associated with poor prognosis (Jo *et al*. 2014; Ruan *et al*. 2017; Aguilar-Medina *et al*. 2019). In the normal endometrium, SOX9 labels putative epithelial stem/progenitor cells and is thought to regenerate the epithelium (Cousins *et al*. 2021). Furthermore, the expression of SOX9 is significantly higher in endometrial cancers and atypical proliferative lesions (Saegusa *et al*. 2012; Gonzalez *et al*. 2016). Intriguingly, transgenic overexpression of SOX9 causes endometrial hyperplasia (Gonzalez *et al*. 2016). Herein, we observed that in the controls, SOX9 labeled some cells of the glandular epithelium in the basalis region. In young hypomorphs, many cells of the glandular epithelium were labeled positive for SOX9 and this number increased further as the animal aged. Moreover, as compared to the young hypomorphs, the number of cells with nuclear SOX9 increased in the aged hypomorphs. These observations suggested that the expansion of epithelial cells in absence of HOXA10 appears to be that of the SOX9 positive progenitor cells. Although more markers need to be investigated to prove this point. Since HOXA10 is required for cell differentiation in the endometrium (Ashary *et al*. 2020), it is possible that in absence of HOXA10, the epithelial cell progenitors do not differentiate correctly and continue to expand leading to hyperplasia which eventually progresses to cancer. Also, SOX9 and Wnt-β catenin pathways cooperate to drive cell proliferation and cancer progression (Santos *et al*. 2016). Elevation of SOX9 and β-catenin levels upon loss of HOXA10 provide substantial evidence to suggest that HOXA10 is upstream to Wnt-β catenin and SOX9, and activation of these networks could be one of the prime reasons for endometrial hyperplasia and cancer phenotypes.

Studies have established that the hippo signaling pathway mainly its downstream effector YAP1 functions as an oncogene in most cancers (Mo *et al*. 2014; Moroishi *et al*. 2015). In endometrial cancers, YAP1 expression is elevated, and knocking down YAP1 inhibits cell proliferation while its overexpression promotes proliferation, anchorage-independent growth, invasion, migration, and chemoresistance of endometrial cancer cell lines (Rosenbluh *et al*. 2012; Tsujiura *et al*. 2014). Herein, we observed that in the young HOXA10 hypomorphs, YAP1 was elevated and the localization was mainly cytoplasmic with few cells having nuclear expression. However, in the aged animals, YAP1 expression was significantly elevated and it was mainly nuclear. In most cancers, YAP1 and β-catenin cooperatively bind to the promoters of anti-apoptotic genes and suppress their expression thereby driving tumorigenesis (Rosenbluh *et al*. 2012). The pro-survival effects of endometrial epithelial cells and their progression to cancer in a low HOXA10 environment could be due to the overactive Hippo and Wnt signaling pathways. Further, studies using specific inhibitors will be required to delineate the mechanistic basis of this hypothesis.

In summary, we demonstrate that HOXA10 plays a crucial role in the development of endometrial hyperplasia which progresses to well-differentiated endometrial adenocarcinoma with age. This progression is coupled with increased expression of members of Wnt-β catenin pathways, SOX9, and YAP1. For the first time, we demonstrate a clear role of HOXA10 in the development and progression of endometrial cancer and suggest a framework for developing anti-HOX therapies for the control and treatment of endometrial cancers. Thus, further studies are needed to fully illustrate the mechanisms by which HOX genes contribute to cancer predisposition and progression to eventually translate these findings into the development of new strategies for precision cancer medicine.

## Declaration of interest

The authors declare that no conflict of interest that could be perceived as prejudicing the impartiality of the research reported.

## Funding

The manuscript bears the NIRRH ID: RA/1234/03-2022. The study was funded by grants from the Department of Science and Technology (SERB) (SR/SO/HS-0277/2012), Govt of India to DM. AM is the recipient of the senior research fellowship from ICMR and University Grants Commission (UGC), Govt of India (# 2121330598). DM lab is funded by grants from ICMR, Govt of India. SSM acknowledges funding from JC Bose Fellowship (SERB-JCB/2017/000027)

## Author contribution

AM: Planned and performed the experiments, data acquisition, data analysis, and wrote the manuscript

NG: Experimental data collection, and primary analysis

SSM: Experimental inputs, primary analysis, and manuscript corrections

DM: Conceptualized the work, planned the project and wrote the manuscript

## Acknowledgments

We thank the staff of the Animal House (NIRRCH) for their help during the animal maintenance

## Notes

### Competing Interest Statement

The authors have declared no competing interest.

