## Supplementary material for "Loss of HOXA10 causes endometrial hyperplasia progressing to endometrial cancer": Supplymentery Tables

**Supplementary Table1**: Primers used in the study; sequence, annealing temperature and product size

| **Gene** | **Sequence** | **Annealing Temperature (⁰C)** | **Product Size (bp)** |
| --- | --- | --- | --- |
| ***Genotyping primer*** | F-5' ACGTCGAGGTGCCCGAAGGA 3'; R-5’ ACGCTATGTGGATACGCTGCTT 3’ | 68⁰C | 543bp |
| ***Ires*** | F-5' CCTCGGTGCACATGCTTTAC 3'; R-5’ CGAGTACAAGCCCACGGTG 3’ | 62 ⁰C | 130bp |
| ***Hoxa10*** | F-5'ACTAAGAGCAGCACGGTACG3'; R-5'CCTTTGGAACTGCCTTGACTC3' | 58⁰C | 200bp |
| ***Esr1*** | F-5' CATAACAGCCTCGGAACGGA 3'; R-5’ GGGCCACCTGCTTGAGAAGA 3’ | 63⁰C | 156bp |
| ***Esr2*** | F-5' TGGCTGGGCCAAGAAAATCC 3'; R-5’ CCTCATCCCTGTCCAGAACG 3’ | 65⁰C | 170bp |
| ***Wnt4*** | F-5’TGGACTCCCTCCCTGTCTTTGGGA3’  R-5’TCCTGACCACTGGAAGCCCTGTG3’ | 64⁰C | 188bp |
| ***Ctnnb1*** | F-5’GCGGCCGCGAGGTACCTGAA3’  R-5’GAAGGAGCTGTGGTGGTGGCA3’ | 60⁰C | 192bp |

**Supplementary Table 2:** The primary antibody and their concentration used for immunohistochemistry

| Primary Antibody | Concentration | Source | Catalogue No. |
| --- | --- | --- | --- |
| HOXA10 | 1:250 | Biomatik; Cambridge, Canada | CAE03449 |
| OVGP1 | 1:500 | Abcam; Cambridge, United Kingdom | Ab74544 |
| ERα | 1:30 | Abclonal, Woburn, USA | A3198 |
| ERβ | 1:500 | Novus Biologicals, Colorado, US | NB120-3577 |
| Ki-67 | 1:1500 | Novus Biologicals, Colorado, US | NB110-89717 |
| SOX9 | 1:100 | Novus Biologicals, Colorado, US | NBP1-85551 |
| β-catenin | 1:50 | Cell Signaling, Massachusetts, US | D10A8 |
| YAP1 | 1:2000 | Abcam, Cambridge, UK | ab205270 |
