## Supplementary figures and images for "Loss of HOXA10 causes endometrial hyperplasia progressing to endometrial cancer"

### Supplementary Fig 1

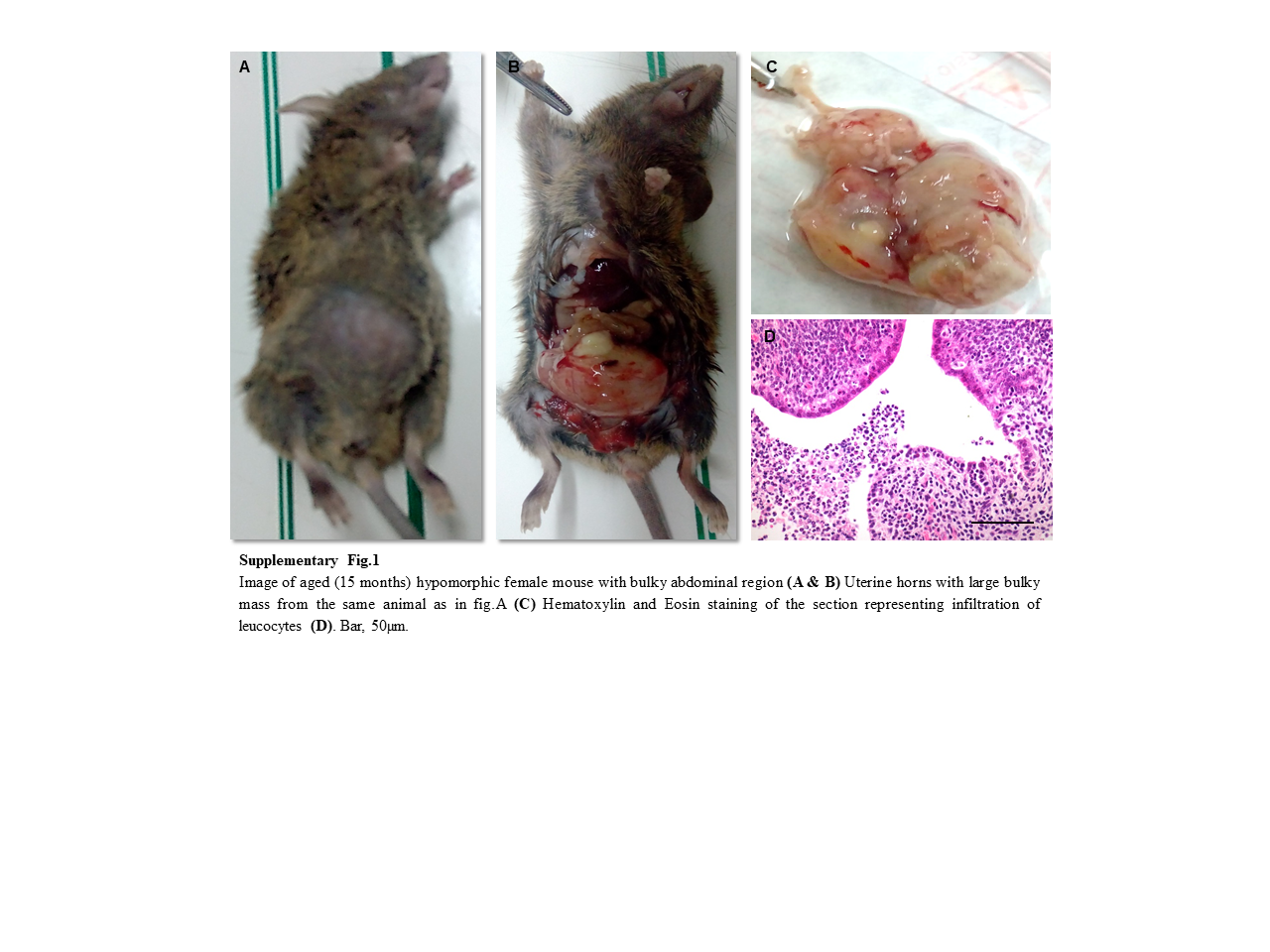

### Supplementary Fig 2

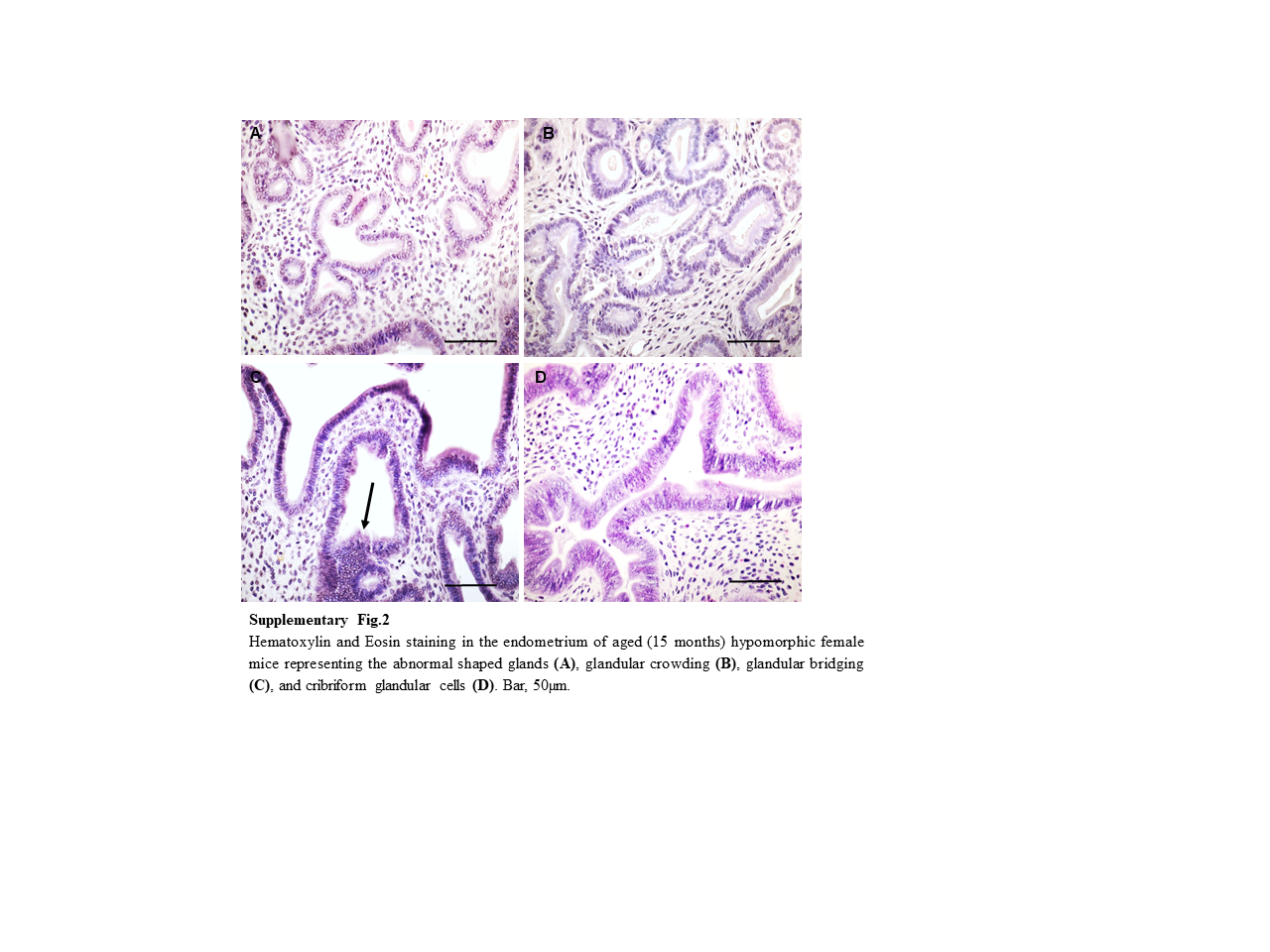
